# Five novel ubiquitous totiviruses function as virulence-promoting symbionts in the obligate biotrophic fungus *Puccinia triticina*

**DOI:** 10.64898/2026.08.25.747185

**Authors:** Jinyang Li, Zhiwen Zheng, Nuoheng Wang, Huiguang Zhao, Yanan Lu, Na Liu, Pengyu Song, Zhenling Ma, Wenming Zheng, Yanhui Zhang

## Abstract

Mycoviruses modulate fungal fitness and pathogenicity, yet their biological roles in obligate biotrophic phytopathogens remain poorly understood. Here, we report the first functional characterization of totiviruses in rust fungi, identifying five novel totiviruses, designated *Puccinia triticina* totivirus 1 to 5 (PtTV1–PtTV5), from the wheat leaf rust fungus *Puccinia triticina* (*Pt*). All five PtTVs possess the canonical genomic architecture of *Totiviridae*, including two overlapping ORFs and a conserved−1 ribosomal frameshifting motif. Transcriptional profiling revealed that PtTVs are highly expressed during early Pt infection. PtTV-encoded proteins suppressed BAX-triggered programmed cell death in *Nicotiana benthamiana*, indicating immune-suppressive activity. Using BSMV-mediated host-induced gene silencing (HIGS), we showed that knockdown of PtTV transcripts significantly impaired fungal hyphal expansion and uredinial formation, concomitant with enhanced host H₂O₂ accumulation. A survey of 90 *Pt* field isolates from four major wheat-growing regions of China revealed that PtTVs are ubiquitously distributed in natural rust populations. Collectively, these findings demonstrate that totiviruses function as virulence-promoting symbionts in *Pt*, establishing for the first time a functional link between totiviral infection and enhanced pathogenicity in cereal rust fungi and identifying candidate targets for RNAi-based disease control.

**IMPORTANCE:** Obligate biotrophic rust fungi represent a major group of plant pathogens with severe global agricultural impacts, yet the biological functions of their native mycoviruses remain largely uncharacterized. This study identifies five novel totiviruses from the wheat leaf rust pathogen *Pt* and provides the first functional demonstration that totiviruses act as virulence-promoting symbionts in cereal rust fungi. The ubiquitous presence of these viruses across geographically diverse field isolates highlights their broad ecological relevance in natural pathogen populations. These findings advance our understanding of fungus-virus mutualism in obligate biotrophic systems and offer a promising, durable molecular target for RNA-based sustainable management of wheat rust diseases.

## INTRODUCTION

Mycoviruses are ubiquitous symbionts across all major fungal lineages, forming persistent intracellular associations that shape host fitness, stress tolerance, and pathogenicity(1, 2). While most mycoviral infections are asymptomatic, two divergent symbiotic outcomes have been well documented. Hypovirulence-inducing mycoviruses attenuate pathogen aggressiveness and have been extensively explored as biocontrol agents, as exemplified by Sclerotinia sclerotiorum hypovirulence-associated DNA virus 1 (SsHADV-1)(3, 4). In contrast, some mycoviruses act as mutualistic partners to enhance fungal virulence, an evolutionary strategy that supports stable vertical transmission within host populations(5).

The family *Totiviridae*, a major mycoviral lineage, includes non-enveloped icosahedral *Totivirus* members that predominantly infect fungi and carry a single linear dsRNA genome of 4.6-7.0 kb, with two overlapping open reading frames (ORFs) encoding the capsid protein (Gag/CP) and the RNA-dependent RNA polymerase (RdRp/Pol)(3, 6–8). A Gag-RdRp fusion protein is typically generated via a conserved −1 ribosomal frameshifting mechanism(9, 10). Totiviruses replicate exclusively in the host cytoplasm and generally establish long-term commensal or mutualistic relationships with their fungal hosts(10). The best-characterized system is Saccharomyces cerevisiae virus L-A (ScV-L-A), which confers a killer toxin phenotype and enhances host competitive fitness, representing a classic paradigm of fungus–virus mutualism(11). However, the biological functions of totiviruses in filamentous plant-pathogenic fungi, particularly in obligate biotrophic pathogens that cannot be cultured axenically, remain severely understudied.

Wheat leaf rust, caused by the obligate biotrophic fungus *Pt*, constitutes a persistent global threat to wheat production and food security(12–14). Although mycoviruses were first detected in rust fungi decades ago, functional characterization of rust-associated viruses remains severely limited(15). In the related stripe rust pathogen *Puccinia striiformis* f. sp. *tritici* (*Pst*), totiviruses, mitoviruses, and narnaviruses have been identified at the genomic level, yet only a single narnavirus (Puccinia striiformis virus 5, PsV5) has been functionally linked to virulence modulation(16–19). For *Pt*, recent metatranscriptomic surveys have recovered abundant viral sequences spanning diverse families, including several totivirus-like contigs with preliminary taxonomic classification(20, 21). However, the global diversity of *Pt*-associated totiviruses remains poorly understood, and no mycovirus in *Pt* has been subjected to experimental functional validation. Specifically, whether totiviruses contribute to rust pathogenicity remains a critical unresolved question, and no totivirus has been experimentally demonstrated to enhance virulence in any cereal rust pathogen.

To address these knowledge gaps, we identified and characterized five novel totiviruses (PtTV1–PtTV5) from *Pt*. By integrating genomic characterization, phylogenetic analysis, heterologous immune assays, BSMV-mediated HIGS, and a nationwide population survey, we provide robust evidence that these ubiquitous totiviruses act as positive regulators of *Pt* virulence. Our study provides the first functional evidence for totivirus-mediated virulence enhancement in cereal rust fungi, and fills a long-standing gap in mycovirus biology in obligate biotrophic pathogens.

## RESULTS

### Genomic organization and molecular characterization of five totiviruses infecting *Pt*

Mycoviruses are pervasive symbionts of plant pathogenic fungi, yet the genomic diversity and molecular characteristics of mycoviruses associated with the wheat leaf rust pathogen *Pt* remain poorly characterized. To fill this knowledge gap, we conducted a systematic survey for dsRNA mycoviruses using urediniospores of *Pt* strain Pt15, which was isolated from naturally infected wheat leaves in China (Fig. 1A). After nucleic acid extraction, novel totivirus-related nucleic acid fragments were obtained, which were resistant to digestion by DNase I and S1 nuclease, suggesting the presence of dsRNA-like nucleic acid molecules (Fig. 1B). Here, we determined the complete genomic sequences of these five putative totiviruses, designated Puccinia triticina totivirus 1 to 5 (PtTV1–PtTV5), via rapid amplification of cDNA ends (RACE). The finished genomes range from 4,829 to 5,111 nucleotides (nt) in length: 4,983 nt for PtTV1, 5,087 nt for PtTV2, 4,829 nt for PtTV3, and 5,111 nt for both PtTV4 and PtTV5. Blastx searches confirmed that all five sequences share significant similarity with recognized members of the *Totiviridae*. In keeping with the canonical genomic organization of totiviruses, each PtTV genome harbors two open reading frames (ORF1 and ORF2), predicted to encode the CP and RdRp, respectively (Fig. 1C). ORF1 is preceded by a 5′ untranslated region (UTR) of 20–74 nt, with deduced CP molecular masses of 99.55, 92.40, 81.43, 92.57, and 94.49 kDa for PtTV1 through PtTV5. ORF2 is followed by a 3′ UTR of 34–60 nt, with corresponding RdRp molecular masses of 97.10, 101.40, 108.16, 102.08, and 101.54 kDa, respectively (Fig. 1C).

**Fig. 1.**
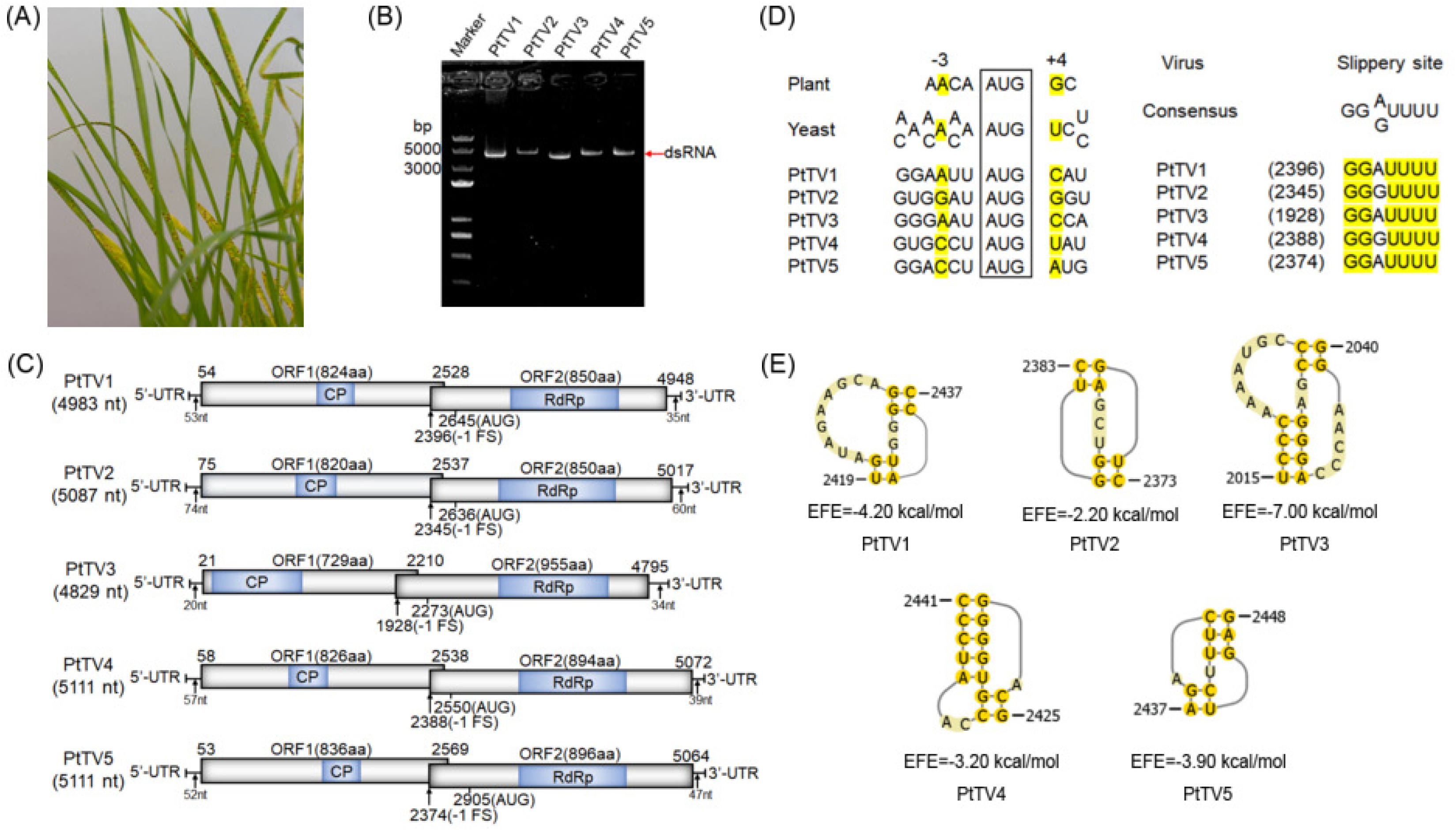
Genome organization and conserved features of PtTV1-PtTV5. (A) Urediniospores morphology of *Puccinia triticina* strain Pt15 grown on wheat cultivar Mingxian 169 for 6 days at 22°C. (B) Agarose gel electrophoresis of dsRNA extracted from urediniospores of *P. triticina* treated with DNase I and S1 nuclease. Marker: DNA ladder 8000 bp. (C) The two overlapping open reading frames (ORFs) and the UTRs are depicted as open boxes and a single line, respectively. Blue shading marks the conserved capsid protein (CP) and RNA-dependent RNA polymerase (RdRp) domains. Black arrows point to the nucleotide coordinates of the ORFs and the predicted −1 frameshift slippery site. (D) The ORF1 start codon (AUG) is boxed (positions +1 to +3). The PtTVs nucleotide sequences at –3 and +4 relative to the AUG start codon (open box, +1∼+3) in plants and yeasts are shown at the top. Putative slippery heptanucleotide motifs (X XXY YYZ) for −1 frameshifting. (E) Predicted pseudoknots downstream of the potential slippery sequences for −1 frameshifting are shown by yellow shadow. Spacer length (nt) and free energy (EFE, kcal/mol) are listed. Numbers (or in parentheses) show nucleotide positions in the contigs.

Examination of the translation initiation contexts showed that the AUG start codons of all PtTV ORFs lie within a Kozak consensus compatible with efficient translation initiation in both fungal and plant cellular environments (Fig. 1D). Sequence analysis of PtTV revealed an overlap region between ORF1 and ORF2. This structural arrangement raises the possibility that ORF2 is translated as a fusion protein with ORF1 through a −1 ribosomal frameshift, which occurs at a typical slipping site ‘XXXYYYZ’ within the overlapping region (where X represents A/C/G/U, Y represents A/U, and Z represents A/C/U)(22). The slippery sequence ‘GGA/GUUUU’ identified in PtTV1-PtTV5 shows marked similarity to that of ScV-L-A (Fig. 1D). For each putative slippery site, a pseudoknot structure was predicted downstream using the DotKnot software, and its minimum free energy was calculated (Fig. 1E). RNA pseudoknot can cause pausing of elongating ribosomes and enhance the efficiency of frameshifting(16, 22). Collectively, these findings strongly indicate that PtTVs employ a conserved −1 PRF mechanism for RdRp expression, mirroring the archetypal gene expression program of prototypical totiviruses.

Conserved Domain Database (CDD) searches and multiple sequence alignment of the predicted RdRp sequences revealed eight canonical conserved motifs (motifs I–VIII), including the diagnostic GDD tripeptide motif that defines the RdRps of *Totiviridae* and other dsRNA viruses (Fig. S1). The presence of these lineage-defining motifs further corroborates the taxonomic placement of PtTV1–PtTV5 within the family *Totiviridae*. The complete genome sequences have been deposited in GenBank under accession numbers PZ417034–PZ417038.

### Phylogenetic analysis confirms five distinct *Totivirus* species with divergent evolutionary histories

To resolve the taxonomic status and evolutionary relationships of the five PtTVs, maximum-likelihood (ML) phylogenetic trees were constructed based on the deduced amino acid sequences of the CP and RdRp, respectively (Table S2). The CP- and RdRp-based trees showed highly congruent topologies (Fig. 2). RdRp, the primary taxonomic marker for double-stranded RNA viruses, was used for fine-scale phylogenetic analysis. Members of the genus *Totivirus* clustered into two well-supported major clades (Group I and Group II), with all five PtTVs falling within Group I. Four viruses (PtTV1, PtTV2, PtTV4, and PtTV5) were assigned to subclade I-D, while PtTV3 occupied a separate phylogenetic position in subclade I-C, indicating clear evolutionary divergence among the five viruses. Pairwise sequence comparison further validated their status as five distinct novel species. PtTV1, PtTV2, PtTV4, and PtTV5 shared only 30.5%–39.84% amino acid identity in both RdRp and CP, and PtTV3 exhibited even deeper sequence divergence from the other four (23.80%–25.54% for RdRp, 11.40%–12.40% for CP), all well below the species demarcation threshold for the genus. Within subclade I-D, PtTV1 shared the highest sequence identity (66.35%) with Uromyces fabae totivirus. PtTV2 and PtTV4 were most closely related to Puccinia striiformis totivirus 3 and Puccinia striiformis totivirus 1 (68.97% and 43.76% identity, respectively), while PtTV5 clustered with Cronartium ribicola totivirus 3 (38.62% identity). PtTV3 was phylogenetically closest to Umbelopsis gibberispora virus 2 in subclade I-C, with 38.28% RdRp amino acid identity (Table S3). Notably, subclade I-D exclusively contains totiviruses isolated from filamentous fungi, consistent with long-term codivergence between these viruses and their fungal hosts. In contrast, subclade I-C harbours viruses from a broader host range including fungi, oomycetes, and insects(23), implying a more dynamic evolutionary history with potential host-shift events for PtTV3. Taken together, phylogenetic and sequence similarity analyses confirm that PtTV1–PtTV5 represent five distinct novel species in the genus *Totivirus* (family *Totiviridae*). Their divergent phylogenetic placements suggest distinct evolutionary origins and potential functional differentiation, which aligns with the divergent temporal expression patterns observed during infection.

**Fig. 2.**
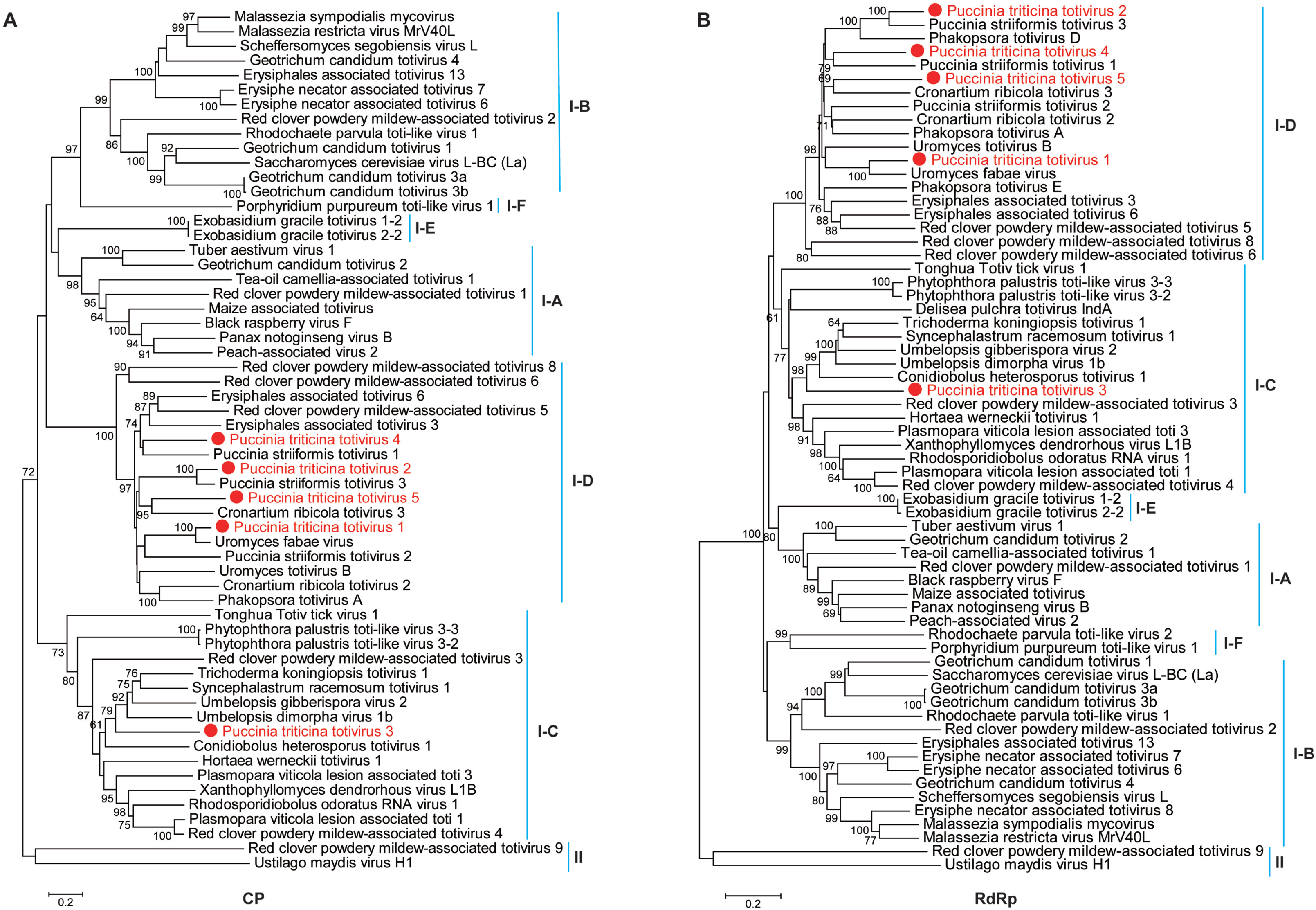
Phylogenetic analysis of PtTVs. Maximum-likelihood (ML) method based on the capsid protein (CP) (A) and RNA-dependent RNA polymerase (RdRp) (B) amino acid sequences of PtTVs and other dsRNA viruses in the genus *Totivirus*, using the Molecular Evolutionary Genetics Analysis (MEGA) software version 11.0. The number above the internal branches indicates bootstrap values estimated with 1,000 replications. Scale bar represents the estimated number of amino acid substitutions per site. Red circles indicate the novel mycoviruses PtTVs in the present study. Table S2 for abbreviations of virus names and viral protein accession numbers used for phylogenetic analysis.

### Transcriptional profiling of PtTVs reveals stage-specific activation during early *Pt* infection

To investigate the potential roles of the five novel PtTVs in *Pt* pathogenicity, we quantified their transcript levels via reverse transcription quantitative PCR (RT-qPCR) across a time course of wheat infection, using resting urediniospores as the baseline control. All five PtTVs were significantly transcriptionally upregulated upon infection initiation. Three viruses (PtTV1, PtTV2, and PtTV5) peaked at 6 hours post inoculation (hpi)—a developmental stage corresponding to urediniospore germination and appressorium formation—and maintained high transcript abundance through 24 hpi, the period of haustorial mother cell differentiation and early host colonization. Notably, PtTV3 and PtTV4 exhibited a delayed expression pattern, with transcript levels continuing to rise from 36 to 48 hpi and reaching their maximum at 48 hpi, coinciding with intercellular hyphal expansion and haustorium development. Transcript levels of all five viruses gradually declined from 48 hpi to 144 hpi, during the late stages of fungal vegetative growth and sporulation (Fig. 3). Collectively, all five totiviruses are transcriptionally activated during the early biotrophic infection phase of *Pt,* with divergent temporal dynamics that likely reflect functional differentiation across infection stages. This early activation pattern is concordant with the impaired early colonization phenotype observed upon PtTV silencing, supporting a key role for these viral symbionts in promoting the initial establishment of rust infection.

**Fig. 3.**
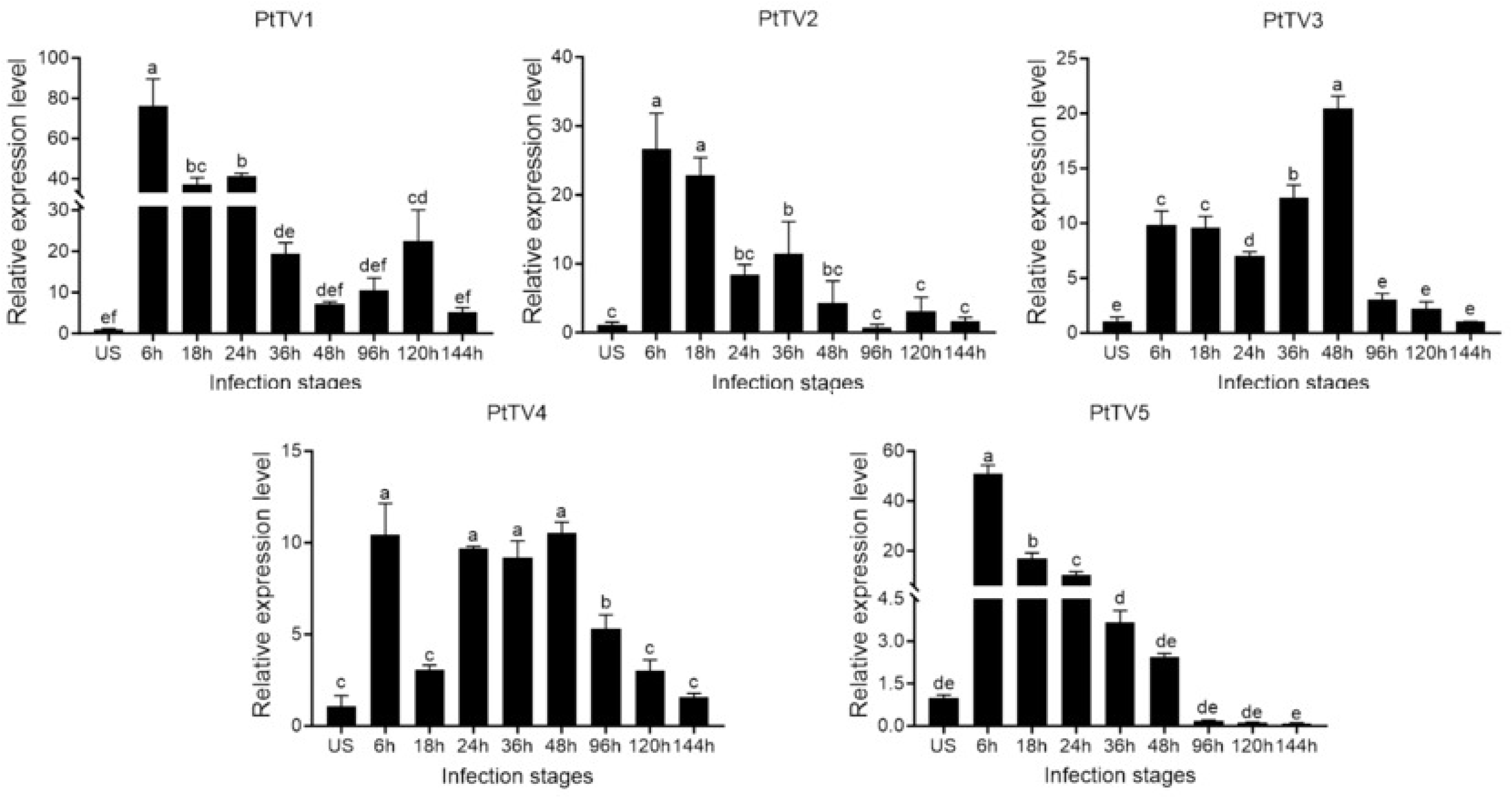
Relative transcript levels of PtTVs at different *Pt* infection stages determined by RT-qPCR. Wheat leaves (wheat cultivar Chinese Spring) inoculated with fresh urediospores (US) (Pt15) were sampled at different time points according to the infection stage of *Pt*. Urediospores were used as a control. Relative transcript level of PtTVs was calculated using the comparative 2^−ΔΔCT^ method. *Actin* was used as the reference gene. Mean and standard deviation were calculated with data from three independent biological replicates. Different letters on the top of each column are significantly different at the P < 0.05 level of confidence according to Tukey’s HSD post-hoc test.

### PtTV-encoded proteins suppress BAX-triggered programmed cell death

Transient expression of the pro-apoptotic Bcl-2 family protein BAX in *Nicotiana benthamiana* triggers programmed cell death (PCD), which recapitulates the hypersensitive response, a hallmark of plant innate immunity(24). To investigate whether PtTVs can interfere with plant immune signaling, we performed *Agrobacterium tumefaciens*-mediated transient expression assays in *N. benthamiana* leaves to test whether PtTV-encoded proteins inhibit BAX-induced PCD. The *Pst* effector PstGSRE1, a well-characterized suppressor of BAX-triggered PCD, was employed as a positive control(25), and an empty vector construct was used as the negative control. At 4 days post infiltration, no necrotic lesions were observed in leaf regions expressing either PstGSRE1 or individual PtTV proteins, whereas clear cell death occurred in the negative control regions (Fig. S2). These results indicate that proteins encoded by PtTVs are capable of suppressing BAX-mediated PCD in a heterologous plant expression system. This immune-suppressive activity is mechanistically consistent with the enhanced host H₂O₂ burst and attenuated fungal virulence observed upon PtTV silencing, supporting a model in which PtTVs promote rust infection by attenuating host innate immune responses.

### Silencing of PtTVs via HIGS significantly attenuates the virulence of *Pt*

To functionally validate the regulatory roles of PtTVs in *Pt* pathogenicity, we performed loss-of-function assays using a BSMV-mediated HIGS system. At 14 days post BSMV inoculation, conspicuous photobleaching was observed in wheat plants infiltrated with BSMV:TaPDS-as, confirming the efficacy and reliability of the HIGS system (Fig. 4A). Mild chlorotic mosaic symptoms appeared uniformly on all BSMV-inoculated leaves, indicating successful systemic viral infection (Fig. 4A). The fourth leaves of BSMV-treated plants were then inoculated with fresh urediniospores of *Pt* strain Pt15. At 7 days post inoculation (dpi), plants inoculated with BSMV:PtTVs constructs exhibited a markedly reduced number of uredinia per leaf compared with the BSMV:γ empty vector control and the mock-inoculated control (Fig. 4B).

**Fig. 4.**
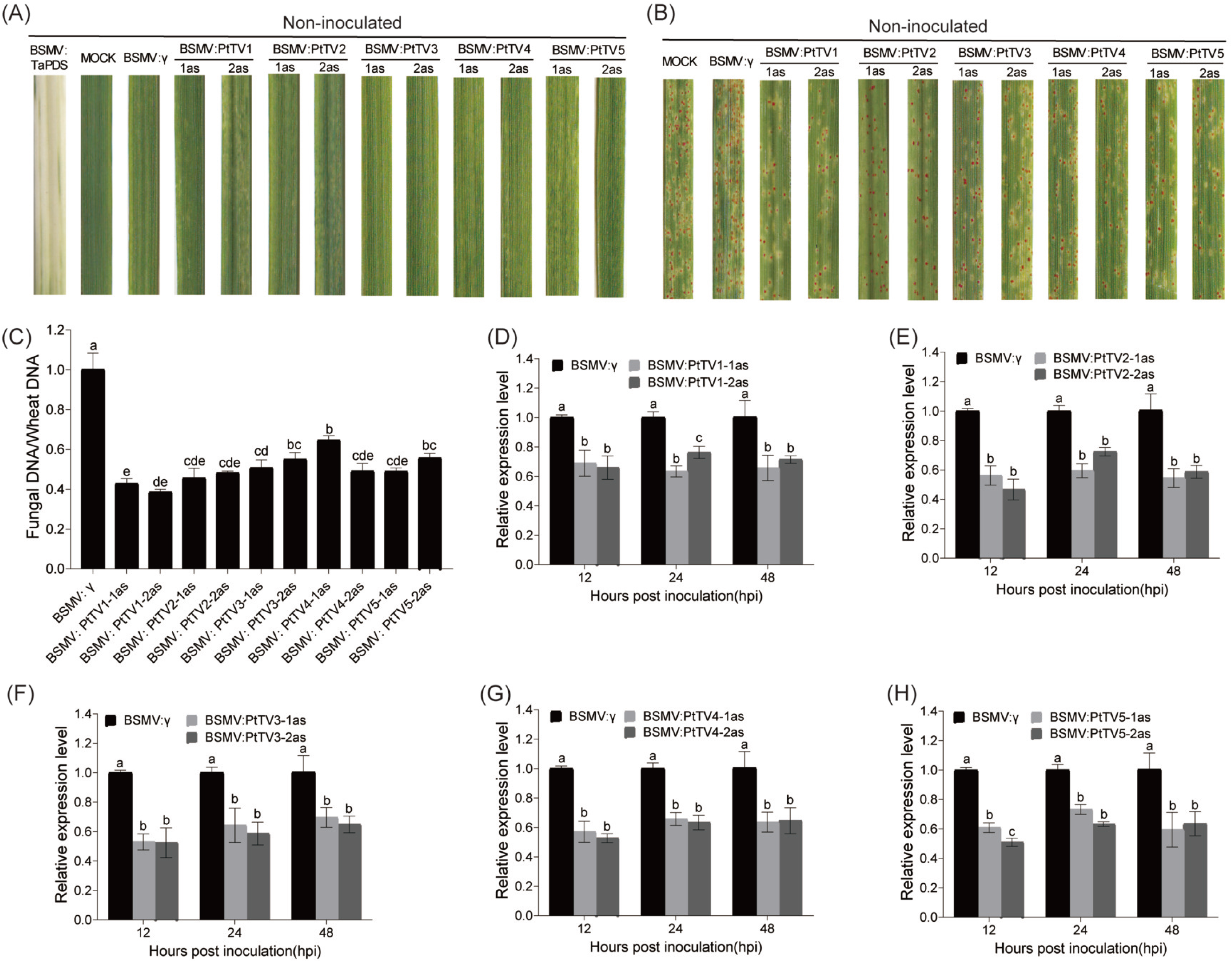
Functional assessment of the role of PtTVs in *Pt* virulence using BSMV-mediated host-induced silencing. (A) Mild chlorotic mosaic symptoms were observed on leaves inoculated with BSMV:γ, BSMV:*TaPDS-as*, or BSMV:PtTVs-1/2as at 14 days post inoculation (dpi) with barley stripe mosaic virus, whereas mock-treated leaves (1× FES buffer) remained asymptomatic. (B) Phenotypes of the fourth leaves of knockdown and control plants at 7 dpi with Pt15. Mock-inoculated and BSMV:γ-treated plants served as negative controls. (C) Fungal biomass was assessed by qPCR. The ratio of fungal to wheat nuclear DNA content was determined using the fungal *Actin* and wheat *TaEF1* genes, respectively. Genomic DNA was extracted from the fourth leaves of three independent plants at 5 dpi with Pt15. (D-H) Relative transcript levels of PtTVs in *Pt*-inoculated leaves were measured by RT-qPCR at 12, 24, and 48 hours post inoculation (hpi). Transcript levels were normalized to *Actin*. Data are presented as means ± standard error (SE) of three independent biological replicates. In panels C-H, different letters above the columns indicate significant differences at P < 0.05 (Tukey’s HSD post-hoc test).

To quantify fungal proliferation, total genomic DNA was extracted from *Pt*-infected leaves, and the relative abundance of fungal *Actin* and wheat *TaEF1* was determined by quantitative PCR (qPCR). The *Pt* biomass ratio (*Actin*/*TaEF1*) was calculated to normalize fungal load against plant tissue. At 5 dpi, all five PtTV-silenced groups showed significantly reduced fungal biomass relative to the BSMV:γ control. For BSMV:PtTV1-1/2as, PtTV2-1/2as, PtTV3-1/2as, PtTV4-1/2as, and PtTV5-1/2as treatments, biomass was reduced to 57%–61%, 52%–54%, 45%–49%, 35%–51%, and 44%–51% of the control level, respectively (Fig. 4C). Importantly, two independent silencing fragments targeting each virus yielded highly consistent results, minimizing the possibility of off-target effects. We further assessed silencing efficiency by measuring the relative transcript levels of each PtTV via RT-qPCR at 12, 24, and 48 hpi. Compared with the BSMV:γ control, BSMV:PtTV1-1/2as reduced PtTV1 transcript levels by 31%–34%, 24%–37%, and 29%–35% at the three time points, respectively (Fig. 4D). Similarly, BSMV:PtTV2-1/2as reduced PtTV2 levels by 44%– 53%, 28%–41%, and 42%–46%; BSMV:PtTV3-1/2as reduced PtTV3 levels by 47%–48%, 36%–41%, and 31%–35%; BSMV:PtTV4-1/2as reduced PtTV4 levels by 34%– 37%, 36%–37%, and 43%–47%; and BSMV:PtTV5-1/2as reduced PtTV5 levels by 39%–49%, 27%–37%, and 37%–41% at the corresponding time points (Fig. 4E through H).

Collectively, these results demonstrate that BSMV-mediated HIGS effectively suppresses the expression of all five PtTVs, and that reduced viral accumulation consistently leads to attenuated *Pt* pathogenicity and restricted fungal growth in wheat. The congruent phenotypes across all five totiviruses further support a conserved virulence-promoting function for this viral group in cereal rust fungi.

### Silencing of totiviruses inhibits hyphal growth and promotes ROS accumulation of *Pt*

To dissect the histological basis underlying the attenuated virulence caused by PtTV silencing, we quantified fungal hyphal length and infection area at 12, 24, and 48 hpi across PtTV-silenced and control plants, using DP-BSW software for image-based analysis (Fig. 5A). At 12 hpi, a developmental stage corresponding to germ tube elongation and haustorial mother cell formation, all five PtTV-silenced treatments exhibited significantly reduced hyphal length compared with the BSMV:γ control, while infection area remained unchanged for PtTV4 and PtTV5. By 24 and 48 hpi, both hyphal length and infection area were significantly decreased across all five silencing groups (Fig. 5B and C). This time-dependent phenotypic defect is concordant with the high transcriptional activity of PtTVs during early infection, suggesting that these viruses predominantly function to support initial fungal colonization and subsequent hyphal expansion in host tissue.

**Fig. 5.**
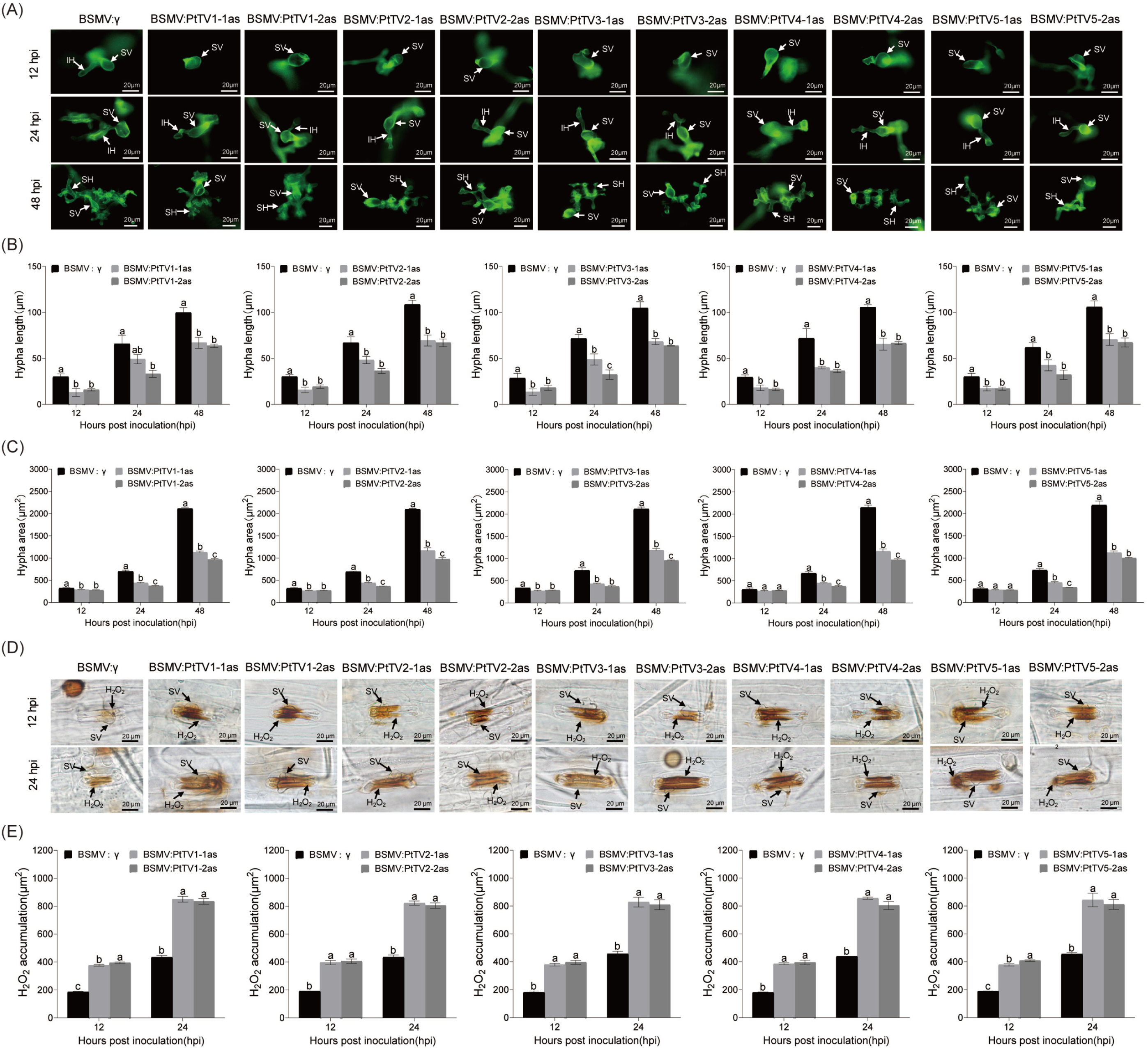
Silencing of PtTVs reduces infectious hyphae growth of *Pt* and increases ROS accumulation in wheat leaves. (A) At 12, 24, and 48 hours post inoculation (hpi), wheat leaves infected with Pt15 were stained with wheat germ agglutinin (WGA), and fungal structures were visualized under a fluorescence microscope. Abbreviations: IH, infection hyphae; SH, secondary hyphae; SV, substomatal vesicle. Scale bar = 20 μm. (B and C) Hyphal length (B) and infection area (C) were measured at the indicated time points. Data were collected from 50 infection sites per treatment across three independent biological replicates. (D) Wheat leaves pretreated with BSMV:γ or BSMV:PtTVs-1/2as were inoculated with Pt15. At 12 and 24 hours post inoculation (hpi), H₂O₂ accumulation was visualized by 3,3′-diaminobenzidine (DAB) staining. Fungal structures (SV, substomatal vesicle) are indicated. Scale bar = 20 μm. Data represent 50 infection sites across three independent biological replicates. (E) The DAB-stained area per infection site was measured using DP-BSW software to estimate H₂O₂ levels. Values are shown as means ± standard error (SE). Different letters denote significant differences (Tukey’s HSD post-hoc test, P < 0.05).

Reactive oxygen species (ROS) burst is a hallmark of plant innate immunity and plays a critical role in restricting biotrophic pathogen invasion, with H₂O₂ being a major, stable ROS intermediate(26, 27). We therefore measured H₂O₂ accumulation in wheat leaves as an indicator of host defense activation following *Pt* inoculation (Fig. 5D). Compared with the BSMV:γ control, silencing of PtTVs resulted in significantly elevated H₂O₂ accumulation at both 12 and 24 hpi (Fig. 5E), which is temporally coupled with the impaired hyphal growth phenotype.

Together, these histological data demonstrate that knockdown of PtTVs impairs early fungal colonization and hyphal proliferation, and this growth defect is tightly associated with intensified host oxidative burst. These findings provide cellular-level evidence that PtTVs promote rust infection by suppressing host innate immune responses.

### PtTVs are ubiquitously distributed in natural *Pt* populations across major wheat-growing regions of China

To investigate the prevalence and natural distribution of PtTV1–PtTV5, we performed RT-PCR screening on 90 *Pt* field isolates sampled from six provinces, covering four major ecological wheat-growing zones in China: the Huang-Huai Winter Wheat Region, the Northwest Winter Wheat Region, the Northwest Spring Wheat Region, and the Northern Winter Wheat Region. The panel included 30 isolates from Henan, 20 from Shaanxi, 14 from Gansu, 16 from Xinjiang, 7 from Hebei, and 3 from Inner Mongolia (Table S4), broadly representing the main wheat leaf rust epidemic areas in China.

Notably, all 90 field isolates tested positive for all five PtTVs (Fig. 6), with no natural isolate lacking any of the five viruses. This near-fixation across geographically and ecologically distinct populations demonstrates that PtTV1–PtTV5 are not sporadic, transient infections, but stably maintained, ubiquitous mycoviral symbionts within Pt populations in Chinese wheat-growing regions.

**Fig. 6.**
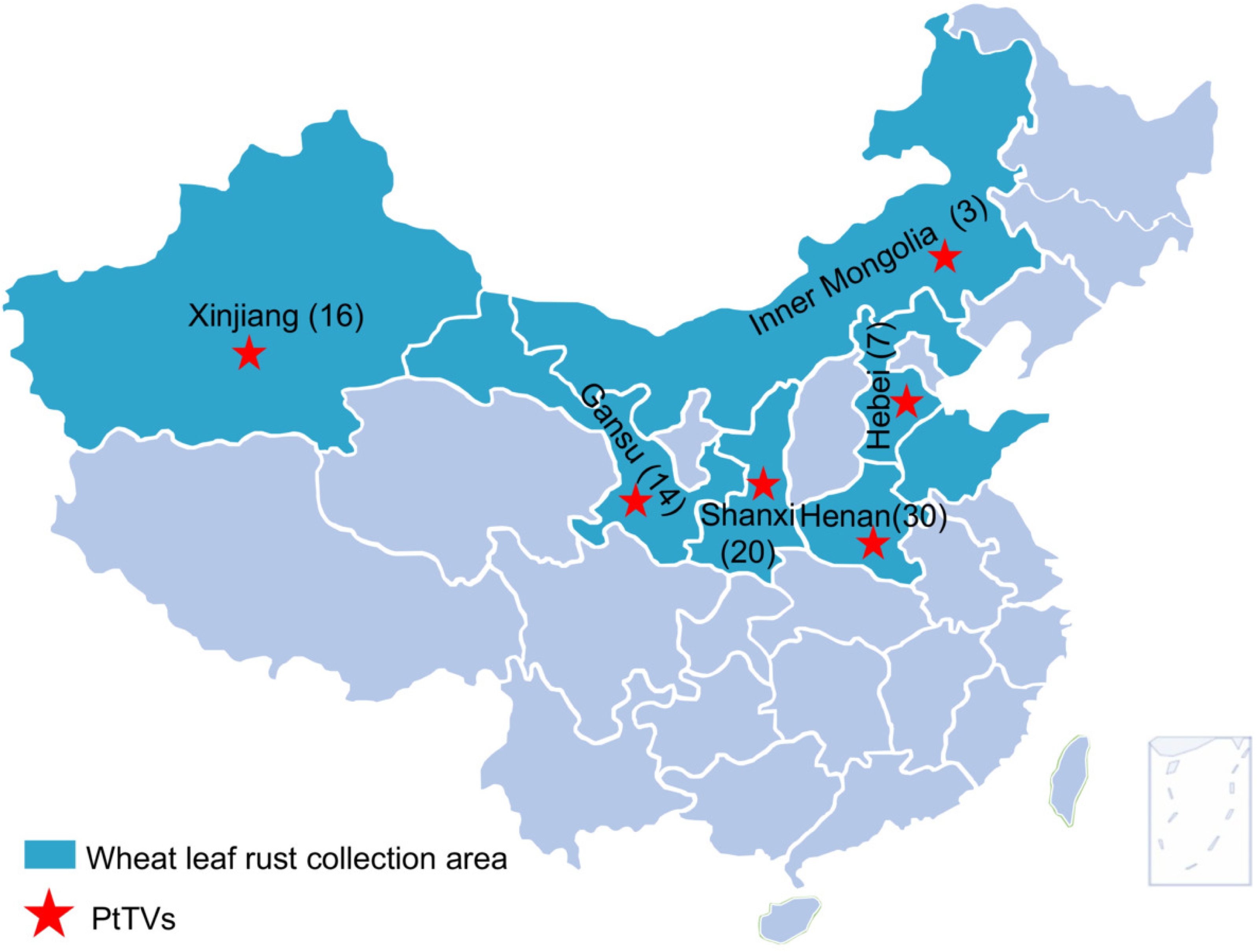
Geographic prevalence of PtTVs. RT-PCR detection was performed on 90 *Pt* isolates collected from six provinces across major wheat-growing regions of China: Henan (30 isolates), Shaanxi (20), Gansu (14), Xinjiang (16), Hebei (7), and Inner Mongolia (3). This sampling scheme allows assessment of the distribution and epidemiological potential of PtTVs in natural *Pt* populations.

**Fig. 7.**
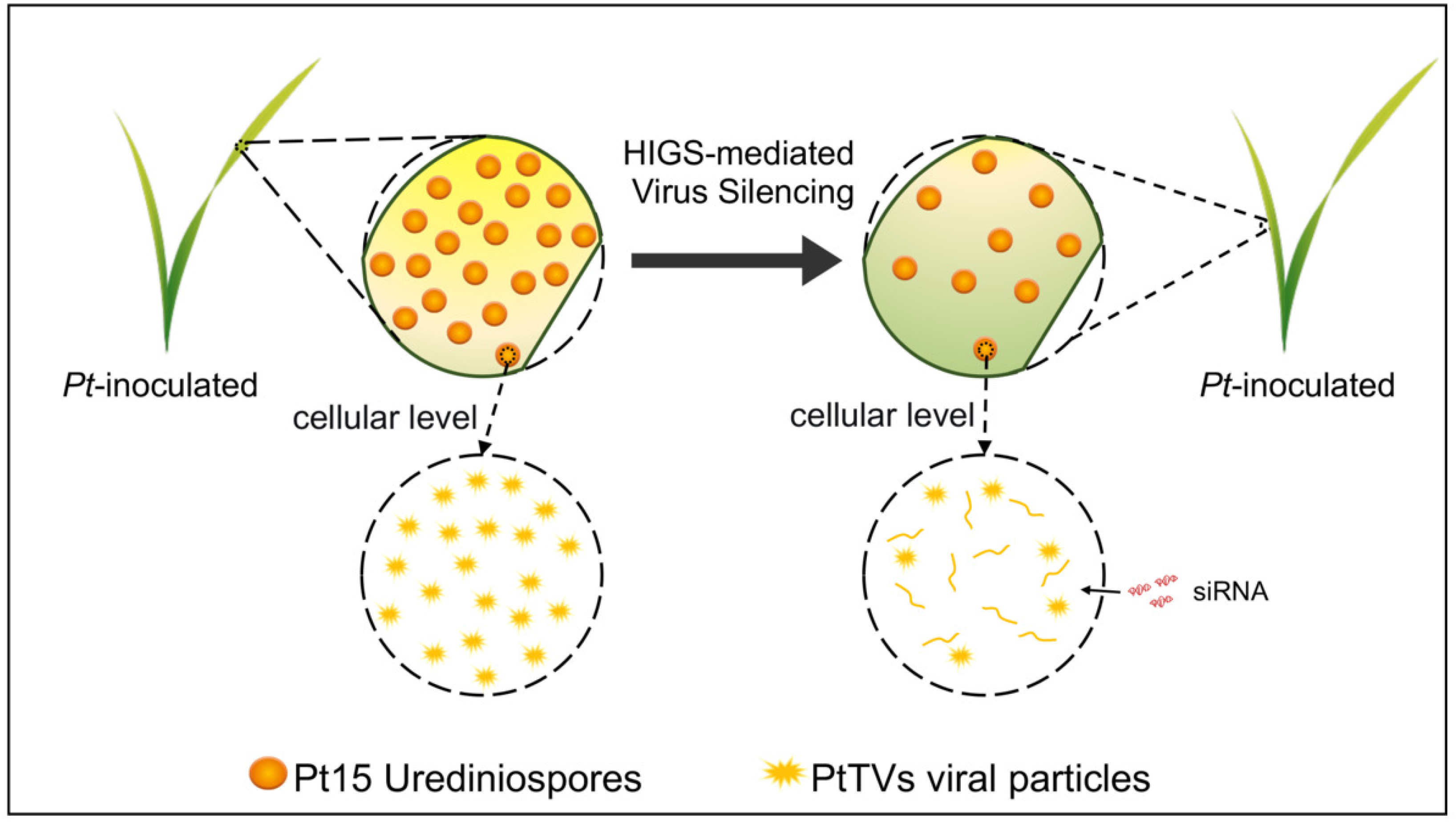
A proposed model for the function of PtTVs during *Pt*-wheat interaction. *Pt* totiviruses function as virulence-promoting symbionts in fungal cells. Host-induced gene silencing (HIGS)-delivered of small interfering RNAs (siRNAs) target and suppress PtTVs accumulation, consequently reducing fungal pathogenicity.

## DISCUSSION

In this study, we identified five novel full-length totiviruses (PtTV1–PtTV5) from the obligate biotrophic wheat leaf rust pathogen *Pt*, and provided the first functional evidence that totiviruses act as virulence-promoting symbionts in a cereal rust fungus. While previous metatranscriptomic surveys have repeatedly recovered fragmented totivirus sequences from rust fungi(16, 20), their biological roles have remained entirely speculative. We combined genomic characterization, heterologous immune assays, HIGS-mediated loss-of-function validation, and nationwide population profiling to establish a robust link between PtTV infection and enhanced rust pathogenicity. This work fills a critical knowledge gap in obligate biotrophic rust virology and expands the paradigm of mycovirus-mediated virulence modulation.

A central question raised by our findings is how totiviruses augment rust virulence at the molecular level. As obligate biotrophs, rust fungi depend entirely on living host cells for nutrient acquisition, and suppression of host programmed cell death and oxidative bursts is a core pathogenic strategy. We show that PtTV-encoded proteins significantly inhibit BAX-mediated cell death in a heterologous plant system, and that silencing of PtTVs elevates host H₂O₂ accumulation and restricts hyphal proliferation in wheat (Fig. 5 and S2). Two non-exclusive mechanisms may underlie this phenotype. First, PtTV infection may transcriptionally or post-transcriptionally modulate the expression of fungal pathogenicity factors, thereby indirectly suppressing host immunity. Second, viral gene products may be translocated into plant cells and act directly as immune suppressors. At present, our data do not distinguish between these possibilities, and the precise molecular pathways through which PtTVs regulate fungal virulence await further investigation. From an evolutionary perspective, this fitness-enhancing symbiosis mirrors the canonical yeast L-A totivirus system, where the virus boosts host competitive ability to ensure its own stable vertical transmission(28–32). A similar virulence-promoting role has also been reported for the narnavirus PsV5 in *Pst*(19), suggesting that mycovirus-mediated virulence enhancement may be a conserved strategy across cereal rust pathogens.

A major technical challenge for studying rust-associated mycoviruses is that obligate biotrophic fungi cannot be cultured axenically or cured of viral infection by conventional methods, which has long hindered functional validation(16). To mitigate this limitation, we have anchored our conclusion on two complementary lines of evidence. First, BSMV-mediated HIGS, a well-established and widely adopted approach for functional gene validation in obligate rust fungi(33), consistently reduced fungal biomass, uredinial formation and hyphal growth across independent assays, coupled with intensified host oxidative burst―all concordant with attenuated virulence upon PtTV knockdown (Fig. 4 and 5). Second, population-level surveillance revealed that PtTV1-PtTV5 are universally present in all 90 field isolates sampled across four major Chinese wheat-growing regions (Fig. 6). This near-fixation in natural populations strongly suggests that PtTVs are not transient infections, but evolutionarily stabilized symbionts that contribute to pathogen fitness under field conditions. Together, genetic and ecological evidence converge to support a virulence-promoting function of PtTVs, even in the absence of isogenic virus-free lines.

More broadly, our findings add to the growing body of evidence that mycovirus-mediated virulence modulation is a widespread, evolutionarily conserved phenomenon across diverse plant-pathogenic fungi. For decades, research emphasis has been placed on hypovirulence-associated viruses as biocontrol agents. Our findings, alongside recent reports of virulence-enhancing mycoviruses in other pathosystems(5, 19), demonstrate that mycoviruses can also function as integral components of the pathogenicity apparatus of plant-pathogenic fungi. This implies that the virome should be systematically incorporated into future studies of rust population evolution, virulence dynamics and disease management. From an applied perspective, the ubiquitous and conserved nature of PtTVs across natural *Pt* populations makes them highly attractive targets for RNA-based disease control. Host-induced and spray-induced gene silencing have proven effective against wheat rust fungi by targeting essential pathogenicity genes(34). Conventional RNAi strategies against rust fungi typically target fungal effector genes, which are prone to rapid sequence diversification and virulence breakdown. In contrast, PtTVs are stably maintained across geographically and phenotypically diverse isolates, implying strong purifying selection and low evolutionary turnover. Targeting these core viral symbionts via HIGS or spray-induced gene silencing could provide broader and more durable protection against wheat leaf rust than effector-targeted approaches.

In summary, this study characterizes five totiviruses from *Pt* and demonstrates for the first time that members of the genus *Totivirus* can positively regulate the virulence of a cereal rust pathogen. The widespread distribution of PtTVs in field populations highlights their ecological relevance and positions them as conserved targets for RNAi-based sustainable control of wheat leaf rust. Future dissection of the viral effector candidates and their host immune targets will further clarify the mechanistic basis of this mutualistic-like symbiosis and inform the design of precise disease intervention strategies.

## MATERIALS AND METHODS

### Fungal strains, plant materials and culture conditions

*Pt* strain Pt15 was isolated from wheat leaf rust fungus in Henan province, China, in 2021. Pt15 was propagated on wheat cultivar Mingxian 169 at 20℃, which is susceptible to *Pt* (Fig. 1A). Fresh urediniospores were collected stored at −80 ℃ for total RNA or viral nucleic acid extraction for further use(35). The wheat cultivar Chinese Spring was used for the analysis of gene expression levels and Fielder was used for BSMV-mediated HIGS assays. The seedlings of wheat and *N. benthamiana* plants were cultured in a controlled environment chamber at 16-22℃ with 16h light/8h darkness. The experiment selected 4 to 6-week-old *N. benthamiana* plants for expression analysis.

### Extraction of dsRNAs, cDNA synthesis and sequencing

The dsRNA was extracted from 1 g of Pt15 urediniospores and using an adsorption column made of cellulose powder CF-11 (Whatman, UK). Isolation of dsRNA was performed in accordance with previously reported procedures with minor modifications(16). The resultant dsRNAs were subsequently treated with DNase I and S1 nuclease (TaKaRa, Dalian, China) to digest genomic DNA and ssRNAs (TaKaRa), followed by quality evaluation via 1.0% (w/v) agarose gel electrophoresis. Using the purified dsRNA fraction as a template, random cDNAs were conducted by RT-PCR(36). The cloning and sequencing of above products were processed using the previously published method(37, 38). Total RNA was isolated from Pt15 and further purified with RQ1 RNase-Free DNase (Promega Corp, Madison, WI, USA) to eliminate fungal DNA. The total RNA was also reverse-transcribed using the RevertAid First Strand cDNA Synthesis Kit (Thermo, MA, USA) for PCR amplification. PCR products were separated on a 1.0% agarose gel and extracted using Hipure DNA Clean up Kit (Magen, Guangzhou, China). The fragments were connected to the pMD18-T (Takara, Dalian, China) vector separately and subjected to Sanger sequencing. The terminal sequences of dsDNAs were determined by RACE. The PCR primers and their sequences mentioned above are listed in Table S1. The complete cDNA sequence was ultimately obtained by splicing together different cDNA clone fragments.

### Sequence and phylogenetic analysis of novel totiviruses

ORFs were predicted using ORF Finder program on the National Center for Biotechnology Information website. Conserved domains were identified using CD-search in the NCBI. Conserved motifs were identified by sequence alignment performed in PROSITE database. To identify RNA pseudoknot structures, prediction was initially performed using DotKnot software, and the predicted structures were subsequently visualized with PseudoViewer3. Multiple alignments of nucleotide and coding amino acid sequences were performed using DNAMAN 7.0 and ClustalW 2.0. For phylogenetic tree construction, nucleotide sequences were imported into MEGA 11.0 software, and subjected to multiple sequence alignment via the ClustalW algorithm. Phylogenetic trees were inferred from the aligned sequences using the maximum likelihood approach with 1000 bootstrap replicates. Phylogenetic trees were separately constructed based on sequence alignments of the putative RdRp and CP regions.

### RT-qPCR analysis

Urediniospores and leaves of wheat Chinese Spring inoculated with Pt15 were harvested at 6, 18, 24, 36, 48, 96, 120, 144 hpi to analyze the expression levels of novel totiviruses. RNA was isolated from all samples with the Quick RNA Isolation Kit (Huayueyang Biotechnology, Beijing, China) according to the manufacturer’s instructions. The RevertAid First Strand cDNA Synthesis Kit (Thermo, MA, USA) was used to generate cDNA, and 2 µL of it as the template were used for PCR amplification.

RT-PCR products were diluted with sterile water at a 1:4 ratio and analyzed by electrophoresis in 1% agarose gel. RT-qPCRwas carried out on the Applied Biosystems StepOne Plus qRT-PCR instrument (ABI, USA), with cDNA mixed with Hieff qPCR SYBR Green Master Mix (Yeasen, Shanghai, China), primers, and distilled water following the manufacturer’s protocol. Primer sequences are listed in Supplementary Table S1. *Actin* were chosen as the internal control genes to normalize the RNA level of *Pt*. Relative gene expression was presented as the expression change of target genes relative to reference genes using the 2^-ΔΔCt^ method, and Statistical significance analysis was performed with Student’s *t*-test. For each sample, three biological replicates and three technical replicates were performed.

### Agrobacterium-mediated transient expression in *N.benthamiana*

The inhibitory activity of the novel totiviruses against BAX-mediated PCD was determined using *A.* infiltration assays. To construct the recombinant plasmid, the RdRp nucleotide sequences of the novel totiviruses were amplified with forward and reverse primers harboring the homologous arms of the potato virus X (PVX) vector pGR106. The amplified fragments were respectively ligated with the target vector using the ClonExpress II One Step Cloning Kit (Vazyme, Nanjing, China). Subsequently, the recombinant plasmids were transformed into GV3101 competent cells, which were cultured in Luria-Bertani medium supplemented with 25 mg·L⁻¹ rifampicin and 50 mg·L⁻¹ kanamycin. The GV3101 strains harboring the corresponding recombinant constructs were resuspended in 10 mM MgCl_2_ solution, adjusted to an OD₆₀₀ of 0.5, and then infiltrated into *N. benthamiana* leaves. After 24 h, *A. tumefaciens* cells carrying the BAX construct were infiltrated into the identical sites of the tobacco leaves. GFP was used as a negative control, and PstGSRE1, which can suppress the proapoptotic protein BAX-induced apoptosis, was employed as a positive control. Symptoms were monitored and recorded at 4 days after infiltration. Each experiment was performed with three independent biological replicates.

### BSMV-mediated gene silencing in wheat

To clarify the potential function of novel totiviruses during the interaction between wheat and *Pt*, two independent fragments ranging from 140 to 200 bp were selected from each viral genome as target sequences for HIGS. Gene silencing primers were designed with Primer3Plus(39), and their specificity was verified by BLASTN alignment against the NCBI database. The cDNA fragments were cloned and subsequently inserted into BSMV following previously established protocol(40). Capped in vitro transcripts derived from the recombinant BSMV genomes, including BSMV:α, BSMV:β, BSMV:γ, BSMV:γ:*TaPDS-as* and the recombinant plasmids BSMV:γ:gene-1as and BSMV:γ:gene-2as, were synthesized using the RiboMAX™ Large-Scale RNA Production System-T7 and the Ribo m⁷G Cap Analog (both from Promega), following the manufacturer’s instructions. The BSMV components BSMV:α and BSMV:β were mixed with either BSMV:γ, or γ:*TaPDS-as*, or a recombinant γ:gene-1as, or γ:gene-2as at a 1:1:1 ratio. The resulting mixture was mechanically inoculated onto the second leaf of two-leaf-stage wheat using 1× FES buffer by gentle rubbing with a gloved finger. For each independent experiment, BSMV:*TaPDS-as* was used as a positive control, while wheat seedlings inoculated with only 1× FES buffer (Sodium pyrophosphate 0.5 g, Bentonite 0.5 g, Kieselguhr 0.5 g, Dipotassium phosphate 2.613 g, Glycine 1.877 g, ddH_2_O to 50 mL) or BSMV:γ were served as a negative control. All BSMV-inoculated wheat seedlings were maintained in controlled environmental growth chambers at 25 ± 2 °C under a 16h light/8h darkness photoperiod. At 14 days after virus inoculation, the fourth wheat leaves were inoculated with fresh Pt15 urediniospores and maintained under a photoperiodic environment with a 16h light/8h darkness cycle at 24 ± 2 ℃. Samples of the fourth leaves were collected at 12, 24, and 48 hpi for RNA extraction that was used for RT-qPCR assay to evaluate the silencing efficiency. Phenotypes of wheat inoculated with Pt15 were identificated and recorded at 7 dpi. The biomass of the fungus and wheat was determined by qPCR-based analysis of genomic DNA. Three independent biological replicates were performed for each experiment.

### Histological observation of *Pt* growth in wheat leaves

The leaves inoculated with Pt15 were taken at 12, 24, and 48 hpi for histological observation as previously described. To visualize the accumulation of hydrogen peroxide (H_2_O_2_), collected wheat leaves were stained with 0.1% 3,3′-diaminobenzidine (DAB) solution. The basal ends of the leaves were immersed in 0.1% DAB buffer and incubated under light for 8-12 h. After sufficient uptake of the DAB solution, the samples were submerged in a destaining solution (Anhydrous ethanol:acetic acid, 1:1, v/v) until the leaf tissues became transparent. Subsequently, the samples were treated with chloral hydrate solution overnight, followed by a Nikon Ni-U microscope (Nikon, Japan) observation and image capture. The area of H_2_O_2_ accumulation was quantified using ImageJ software. The fungus possesses specific structures that enable it to be specifically stained with wheat germ agglutinin (WGA) conjugated to Alexa Fluor 488 (Invitrogen, USA). After destaining, the leaf samples were stained with WGA-Alexa-488.The hyphal length and infected area in the inoculated leaves were then observed and quantified using a Nikon Ni-U fluorescence microscope (Nikon, Japan).

## ACKNOWLEDGMENTS

This study was supported by the Henan Province Major Science and Technology Project of China (221100110100), the National Natural Science Foundation of China (32402328) and the National Key R&D Program of China (2025YFF1000300)

## AUTHOR AFFILIATIONS

^1^State Key Laboratory of Wheat and Maize Crop Science, College of Life Sciences, Henan Agricultural University, Zhengzhou, 450002, China

## AUTHOR CONTRIBUTIONS

Jinyang Li, Formal analysis, Investigation, Methodology, Original draft, Writing – review and editing | Zhiwen Zheng, Methodology, Supervision, Investigation | Nuoheng Wang, Methodology, Investigation | Huiguang Zhao, Methodology| Yanan Lu, Methodology | Na Liu, Resources, Supervision | Pengyu Song, Supervision, Resources | Zhenling Ma, Supervision, Resources | Wenming Zheng, Supervision, Resources, Investigation, Funding acquisition | Yanhui Zhang, Conceptualization, Funding acquisition, Supervision, Writing – review and editing.

## DATA AVAILABILITY

The complete genome sequences of PtTV1–PtTV5 have been deposited in GenBank under accession numbers PZ417034–PZ417038. All other data supporting the findings of this study are available within the article and its Supporting Information files.

## SUPPLEMENTAL MATERIAL

Additional Supporting Information in the Word document.

ASM does not own the copyrights to Supplemental Material that may be linked to, or accessed through, an article. The authors have granted ASM a non-exclusive, world-wide license to publish the Supplemental Material files. Please contact the corresponding author directly for reuse.

